# PNAbind: Structure-based prediction of protein-nucleic acid binding using graph neural networks

**DOI:** 10.1101/2024.02.27.582387

**Authors:** Jared M. Sagendorf, Raktim Mitra, Jiawei Huang, Xiaojiang S. Chen, Remo Rohs

## Abstract

The recognition and binding of nucleic acids (NAs) by proteins depends upon complementary chemical, electrostatic and geometric properties of the protein-NA binding interface. Structural models of protein-NA complexes provide insights into these properties but are scarce relative to models of unbound proteins. We present a deep learning approach for predicting protein-NA binding given the apo structure of a protein (PNAbind). Our method utilizes graph neural networks to encode spatial distributions of physicochemical and geometric properties of the protein molecular surface that are predictive of NA binding. Using global physicochemical encodings, our models predict the overall binding function of a protein and can discriminate between specificity for DNA or RNA binding. We show that such predictions made on protein structures modeled with AlphaFold2 can be used to gain mechanistic understanding of chemical and structural features that determine NA recognition. Using local encodings, our models predict the location of NA binding sites at the level of individual binding residues. Binding site predictions were validated against benchmark datasets, achieving AUROC scores in the range of 0.92-0.95. We applied our models to the HIV-1 restriction factor APOBEC3G and show that our predictions are consistent with experimental RNA binding data.

## Introduction

Protein–nucleic acid (NA) interactions play a critical role in biomolecular processes, including the transcription, translation, regulation, and structural organization of the genome. The ability of NA binding proteins (NABP) to recognize and bind their targets is determined by the spatial arrangement of solvent-exposed side chains and the corresponding geometric, chemical, and electrostatic properties of the protein structure. Structural models of protein-NA complexes offer detailed insights into the physical mechanisms that underly protein-NA recognition but are difficult to obtain experimentally, and accurate *in-silico* structural modeling methods are limited.

Currently, approximately 44,000 protein structures are available in the protein data bank (PDB)^1^ with known NA binding function, with only about 23% of such entries containing complexes of protein bound to NA. In addition to experimentally determined structures, advances in protein structure prediction^2,3^ now offer vast amounts of predicted protein structures that can be directly analyzed with computational methods. It is therefore prudent to develop computational approaches of predicting NA binding which can utilize structural data of NABP in their apo state, determined experimentally or computationally, without the need for models of the protein-NA complex.

In the early days of structural biology, the analysis of experimentally determined structures of protein-NA (PNA) complexes showed that certain physical characteristics of binding interfaces are common among NABP, such as enrichment of polar and positively charged sidechains^4-6^, highly positive electrostatic potential, and the hydrogen bonding proclivity of certain residue– nucleotide pairs^7^. The observation that PNA binding sites contain physicochemical signatures with predictive power led to the development of many computational approaches for predicting NA binding from both protein sequence and structure. Several reviews have been published and comparison studies have been performed which summarize much of this earlier work^8-12^.

Recently, structure–based approaches utilizing graph neural networks (GNNs)^13,14^ have been developed for predicting protein function and binding^15-22^, including NA binding. DeepFRI^15^ is a method for predicting protein function from structure that constructs a graph representation of a protein based on C_*α*_–C_*α*_ atom distances and utilizes a protein-sequence embedding model to generate features for each residue node in the graph. GraphSite^16^ utilizes a similar approach for predicting DNA binding sites, but uses AlphaFold2 for its sequence embedding. GraphBind^17^ is a GNN-based method that represents protein residues as nodes and constructs a dense graph within a sliding sphere of a fixed radius centered on each node. Sequence and structure features are assigned to each node and residue-level DNA and RNA binding site predictions are then learned based on a latent spatial encoding. ScanNet^19^ uses a point-cloud representation of a protein and learns an embedding of the spatio-chemical arrangement of neighboring atoms and residues to predict protein-protein binding interfaces. PeSTo^20^ is a method for predicting protein-protein interfaces using a geometric transformer and only considers atomic coordinates and element type as input information. Their method has been applied to protein-ligand, protein-NA, protein-ion and protein-lipid binding. GeoBind^21^ utilizes quasi-geodesic convolutions over point clouds for DNA and RNA binding site prediction with features closely based on dMaSIF^22^. Other recent methods which do not utilize GNNs but are closely related include MaSIF^23^ which predicts protein-protein binding interfaces and small-ligand binding sites and PST-PRNA^24^ which predicts RNA bind sites.

We present PNAbind, a GNN-based method for predicting DNA and RNA binding function and binding sites from protein structure. In our approach a protein structure is represented using a mesh discretization of the solvent excluded molecular surface. The molecular surface is a convenient representation because it accurately captures important geometrical characteristics of a protein that are relevant for binding, such as binding pockets and shape complimentary of the binding site with a binding target, in a much more efficient way compared to voxel or density-based representations. In addition to the mesh geometry, geometric, electrostatic, and chemical properties of the protein structure are mapped to the surface and included as features. Evolutionary information in the form of multiple sequence alignments can also be incorporated and improve binding site prediction. Our GNN models learn encodings of the spatial arrangement of this information, which are then used for NA binding prediction. We utilize two variants of our networks for two kinds of predictions – local predictions at the level of individual residues for identifying regions of a protein surface that constitutes an NA binding site and global predictions at the level of an entire protein complex for the overall binding function of a protein assembly.

We show that our approach is applicable to proteins with a wide diversity in structural fold, size, and biological function. Our models predict DNA and RNA binding with high accuracy and can discriminate DNA versus RNA binding, which we show is achieved via distinct structural features related to binding mechanisms. Our models for binding site prediction were validated against benchmark datasets and compared with similar recent methods, achieving improvements on a variety of metrics. We show that our method generalizes to both bound and unbound protein structures and produces very low false positive rates on a negative control dataset. Finally, we demonstrate that our model predictions are consistent with experimental RNA binding data for the deoxycytidine deaminase APOBEC3G which plays important roles in the restriction of the HIV-1 virus. Applications of PNAbind include the discovery of unknown NA binding proteins, identification of functional binding residues, better interpretation of biochemical data, and a source of prior information about binding sites for modeling of protein-NA complexes. Data, source code and documentation for PNAbind is available at https://github.com/jaredsagendorf/pnabind.

### PNAbind Overview

PNAbind makes binding predictions based on the geometric, chemical, and electrostatic properties of protein molecular surfaces. Our models are based on GNNs, a versatile neural network architecture designed to operate on graph domains^13,14^, and make inferences using a mesh discretization of the molecular surface. We predict the overall binding function (e.g., DNA-binding or RNA-binding) of a protein or protein assembly via graph classification and predict the location of NA binding sites on the protein surface using graph segmentation. By design, GNNs automatically respect the permutation symmetry of graphs and by choosing vertex and edge features that are invariant under translation and rotation, our models are completely invariant to all isometries of Euclidean space. An overview of our method is shown in Fig. 1, and further details of the network architectures are shown in Fig. 2.

**Fig. 1:**
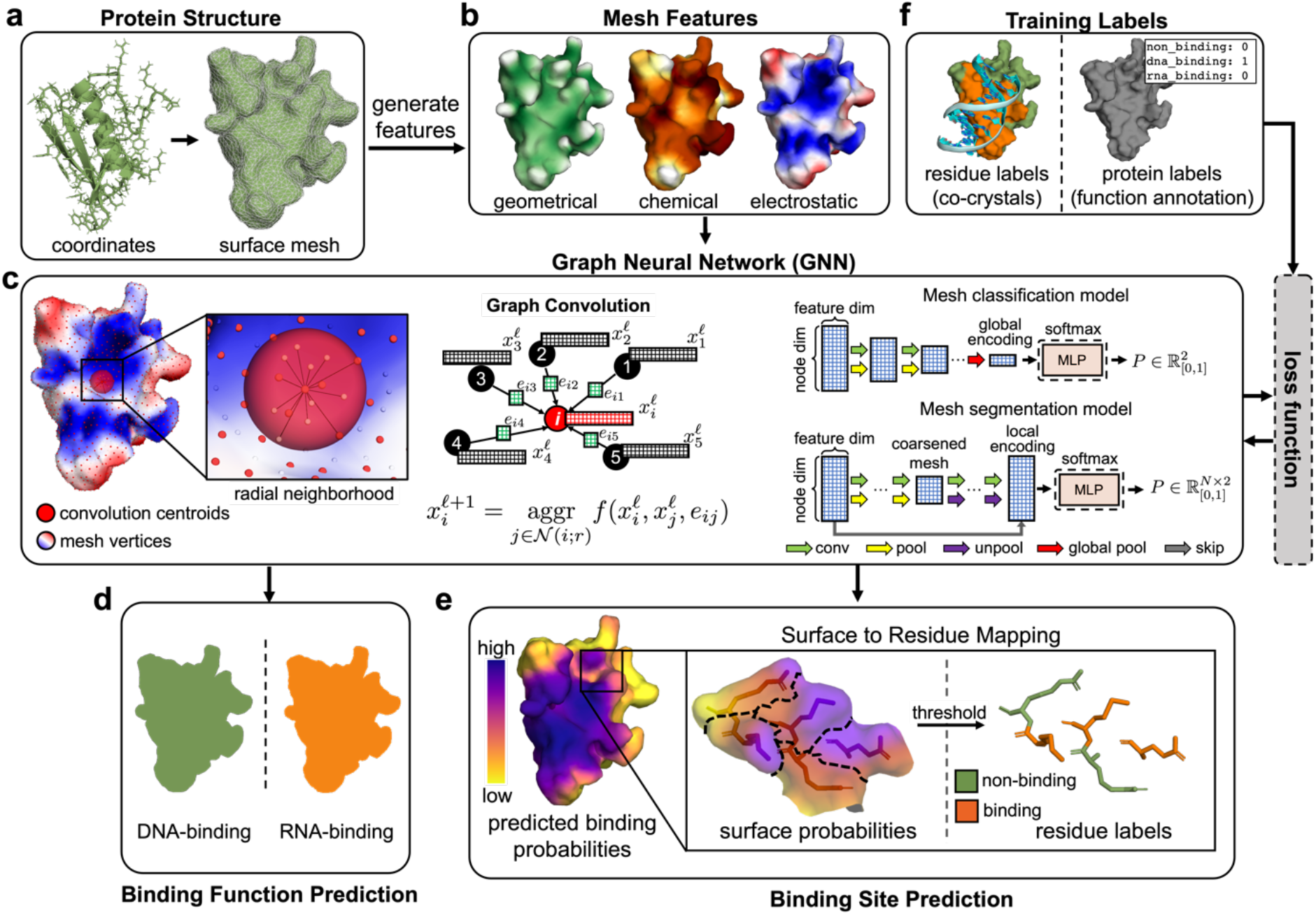
An overview of the PNAbind method. **a** A protein structure is represented as a molecular surface mesh consisting of vertices and edges. **b** Chemical, geometric and electrostatic properties of the protein structure are mapped to the surface mesh. **c** Vertex coordinates are used to compute graph convolutions over radial neighborhoods of the convolution centroids, which are sampled uniformly from the mesh (see Methods). The graph convolution is based on a learned kernel function that is aggregated over edges of the centroid neighbors. Two architectures are used in our method, one for mesh classification (binding function prediction) and one for mesh segmentation (binding site prediction). **d** Output of the classification model is a Bernoulli distribution, representing the probability that the input mesh corresponds to a given functional class (e.g., DNA-binding or RNA-binding). **e** Output of the segmentation model is a probability distribution over the mesh vertices. High probability vertices are identified as NA binding sites, and low probability vertices as non-binding regions. **f** Training data consists of labeled protein surface meshes. Binding site training data is obtained from experimental co-crystal/NMR structures and binding function data is obtained from UniProt functional annotations.

**Fig. 2:**
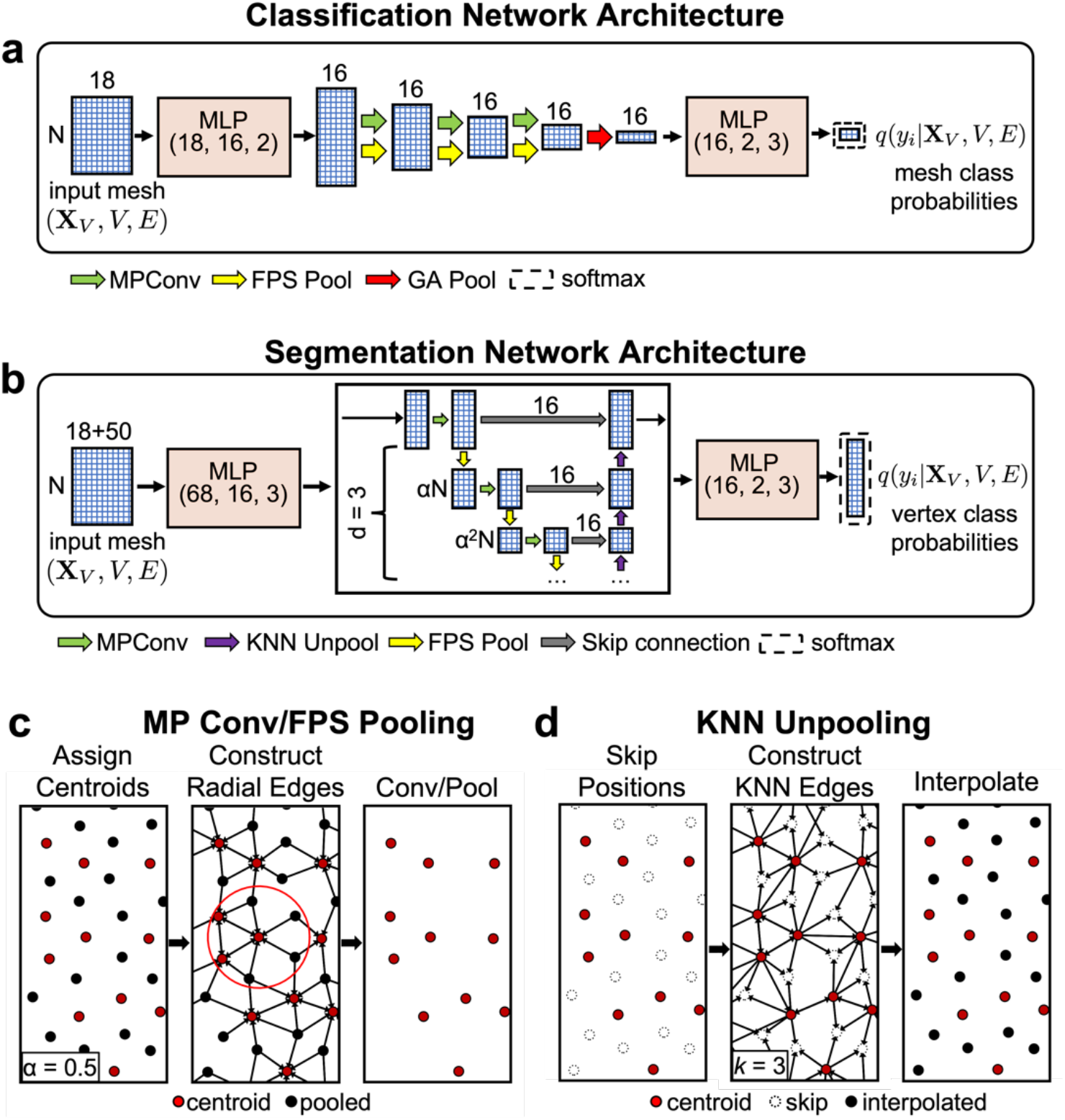
A schematic of model architectures and graph convolution, pooling and unpooling operations. **a** The graph classification network architecture used in our binding function prediction models. The design is analogous to a fully convolutional network. The boxes labeled MLP are fully connected modules (functionally equivalent to 1×1 convolutions) with numbers indicating input size, output size and number of layers. *α* is the sampling ratio used to determine the number of convolution centroids (Methods). The global attention pooling (GA) layer aggregates information over all nodes into a global encoding (Methods). **b** The graph segmentation network architecture used in our binding site prediction models. The design is identical to the classification network up to the GA pooling layer, where instead unpooling layers are used to interpolate the mesh back to full resolution. The overall network topology is that of a graph U-Net^25^ and contains skip connections between pooled and unpooled layers. **c** An overview of the graph convolution and pooling operations used in the networks. First convolution centroids are assigned via farthest-point sampling^26^. Next a radial graph is constructed for all vertices within a fixed radius of any centroid. Graph convolutions are applied over these edges and non-centroid vertices are removed. **d** Unpooling operations are used to restore the mesh to the original resolution for segmentation. For each vertex removed during pooling, its position is stored and a KNN graph is constructed with edges pointing from remaining centroids in the mesh. Feature maps for the restored vertices are then interpolated using distance-weighted averaging over neighboring vertices (Methods).

The primary building blocks of our models are graph convolution, pooling and unpooling layers, followed by multilayer perceptron (MLP) modules (see Methods). Graph convolutions are learned functions applied over the edges of a graph which aggregate information about the local graph structure. A visual example of graph convolutions we use is shown in Fig. 1c. The convolutions are performed over edges defined by a radial distance threshold and the convolution kernel depends on edge features, which describe distance and angle relationships (Fig. 3a) and features of adjacent vertices. By aggregating the activations of the kernel function over the edges of vertex neighborhoods, and applying the convolution repeatedly, spatial arrangements of physicochemical and geometric information that are indicative of NA binding can be encoded. Pooling is used to coarsen the mesh by progressively removing select vertices. It allows for increasing the radius of the next layer of convolutions without incurring a roughly cubic increase in the number of edges in the resulting graph. By increasing the radius, the model can learn encodings over different distance scales. Unpooling is only needed for binding site prediction and it serves to restore the mesh to its original resolution. The final global (local) encodings are then transformed via an MLP module, and the output of the networks represent probability distributions that are computed per-mesh (vertex) for binding function (site) prediction.

**Fig. 3:**
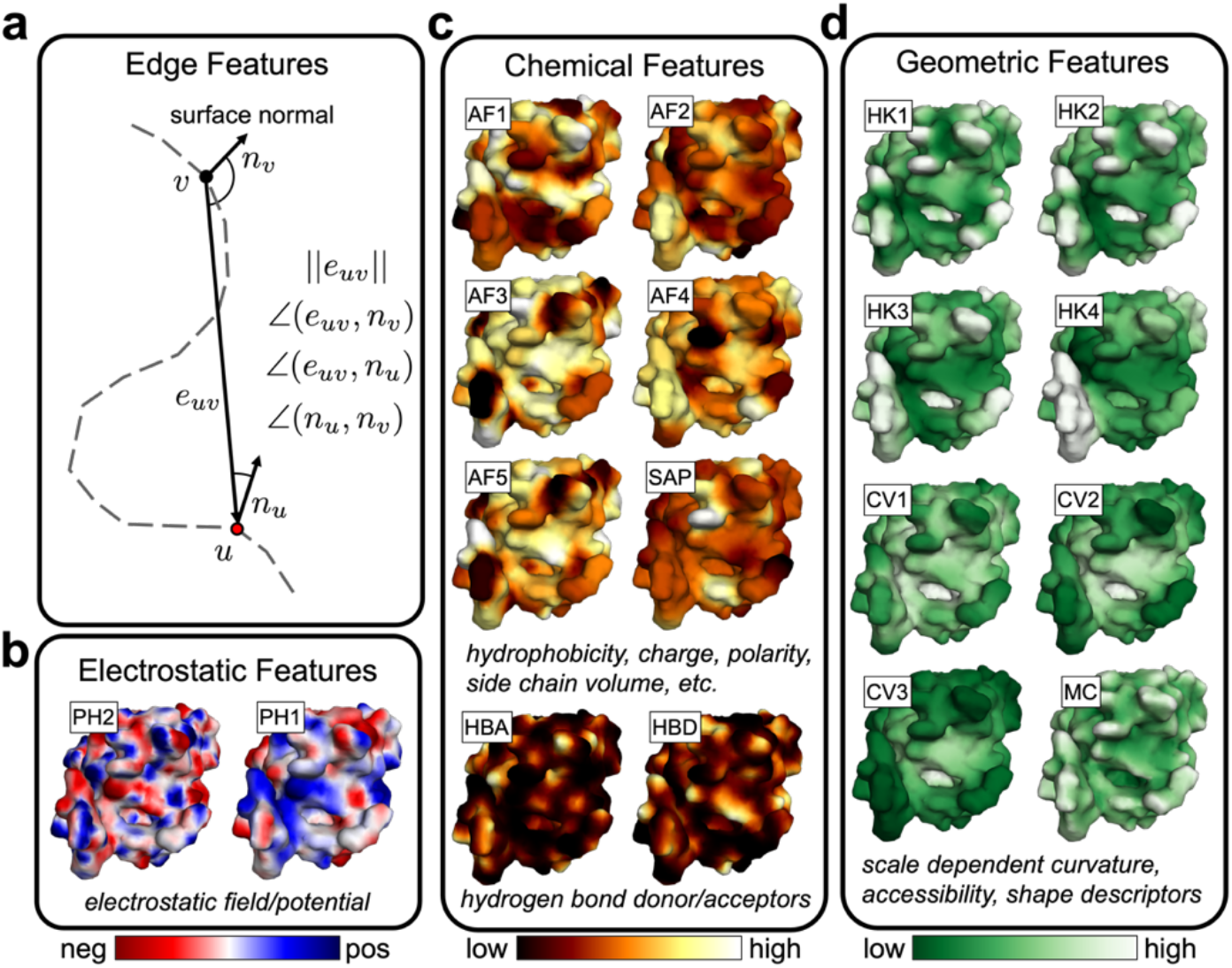
Structure–based edge and vertex features used by PNAbind. **a** Illustrations of edge features. The vectors corresponding to vertex-vertex displacement and surface normals are used to define distance and angle relationships related to the underlying mesh geometry. **b**–**d** Vertex features for the molecular surface mesh corresponding to telomeric single-stranded DNA binding protein Pot1 (PDB ID 4HIM). Structural vertex features are grouped into three categories depending on the type of biophysical properties they describe. Labels for each feature are shown along with brief descriptions. More information about each feature is provided in Methods.

### Surface Mesh Features

We use a combination of features defined on graph edges and vertices that are derived from the atomic resolution protein structure, surface mesh, and sequence in our models. Edge features are used to describe geometric relationships over convolution neighborhoods, such as distance and relative orientation of surface normal vectors (Fig. 3a, Methods). Vertex features encode biophysical properties and evolutionary information that is mapped to the protein surface, which we categorize into four groups. Chemical features describe hydrogen bond donors/acceptors, hydrophobicity, and physical properties of exposed side chains such as secondary structure propensity, charge and polarity. Geometric features are descriptors of the molecular surface and protein structure that depend on its shape and topology. These features can encode distinct shapes that may be related to function such as binding pockets or helical DNA major and minor groove binding elements. Electrostatic features describe the local electrostatic environment of the solvated molecular surface based on the charge distribution and surface geometry. Multiple sequence alignment (MSA) features are based on PSSM matrices computed using PSI-BLAST^27^ and profile HMMs computed using HHblits^28^ (see Methods). In total, 18 structure-based vertex features and four edge features are used in all our models as shown in Fig. 3b–d, and an additional 50 MSA features (20 PSSM + 30 HMM) are used for binding site prediction. Some structural features depend on parameters that define a distance scale for that feature, and different values of these parameters are used to generate multi-scale features that range from high-variance features on the order of ∼3 Å to low-variance features on the order of ∼20 Å (Supplementary Fig. 1).

## Results

First, we report on DNA and RNA binding function prediction using models trained on AlphaFold2^3^ predicted protein structures. Our models predict DNA and RNA binding with AUROC values of 0.94 and 0.95, respectively, and discriminate DNA versus RNA binding with an AUROC of 0.92. Applying attribution methods, we use these models to interpret structural features and regions of the protein structure responsible for binding function, and we highlight chemical and geometric differences that are important for discriminating DNA versus RNA binding. We then show results on NA binding site prediction (residue level predictions) using models trained on native protein structures. Our models achieve high accuracy across a variety of benchmark datasets with AUROC scores ranging from 0.916-0.953. We compare binding site predictions on bound native protein structures versus predicted unbound structures and show high correlations of model predictions. Finally, we apply our model to the deoxycytidine deaminase APOBEC3G and show that our predictions are consistent with what is known experimentally about the RNA binding properties of this protein.

### Prediction of DNA and RNA Binding Function

We developed three datasets for training models to predict DNA and RNA binding function using predicted AlphaFold2 structures available in the AlphaFold database^29^. UniProt^30^ entries with available AlphaFold2 structures were screened based on functional annotations and predicted structure quality and then clustered by sequence similarity to create a DNA binding protein (DBP), RNA binding protein (RBP) and non-binding protein (nBP) dataset (Details and statistics are available in Methods and Supplementary Fig. 4b). Three sets of models were trained – one set to distinguish DBP versus nBP (DnP-6784), the second set to distinguish RBP versus nBP (RnP-6046), and a third set to distinguish DBP vs RBP (RDP-6046). 18 structure-based features were used in these models, (MSA features used for binding site prediction were not included). For each set of models, five-fold cross validation was performed with each holdout fold split into a 50/50 validation/test set and the validation set was used to determine early stopping. Average binary classification metrics on the test splits are shown in Table 1. Our models achieve high accuracy in distinguishing DBP and RBP from nBP with AUROC scores of 0.943 and 0.945, respectively. Our models also distinguish DBP from RBP with high accuracy, achieving 0.920 AUROC despite these proteins binding targets with highly similar chemical structure.

**Table 1:** Binary classification metrics of our models for predicting DNA binding, RNA binding and DNA versus RNA binding function.

| **Dataset Name** | **AUROC** | **AUPRC** | **Recall** | **Precision** | **BA** | **MCC** |
| --- | --- | --- | --- | --- | --- | --- |
| DnP-6784 | 0.943 | 0.940 | 0.896 | 0.893 | 0.872 | 0.746 |
| RnP-6046 | 0.945 | 0.935 | 0.878 | 0.882 | 0.880 | 0.761 |
| RDP-6046 | 0.920 | 0.899 | 0.821 | 0.884 | 0.843 | 0.687 |

**Table 2:** Validation metrics for PNAbind and other recent methods on four test sets used in both our study and the cited studies. Each test set was evaluated using a model trained on the corresponding training set used in the cited studies, namely DNA-573 for DNA-129 and DNA-181, RNA-495 for RNA-117 and RBP09 for RBP61. Metric values for other methods were collected from recent studies^16,17,24^. PNAbind performance is assessed for full biological assemblies and protein monomers, denoted as PNAbind^A^ and PNAbind^M^, respectively.

| RBP61 dataset |  |  |  |  |  |  |  |
| --- | --- | --- | --- | --- | --- | --- | --- |
| Method | AUROC | AUPRC | Precision | Recall | Specificity | MCC | Year |
| PNAbind<sup>A</sup> | **0.951** | **0.875** | **0.786** | **0.823** | **0.924** | **0.735** | 2023 |
| PNAbind<sup>M</sup> | 0.938 | 0.845 | 0.752 | 0.792 | 0.911 | 0.691 |  |
| PST-PRNA<sup>24</sup> | 0.913 | -- | 0.688 | **0.823** | -- | 0.634 | 2022 |
| GraphBind<sup>17</sup> | 0.894 | -- | 0.642 | 0.818 | -- | 0.591 | 2021 |
| aaRNA<sup>34</sup> | 0.862 | -- | 0.587 | 0.782 | -- | 0.528 | 2014 |
| RNA-117 dataset |  |  |  |  |  |  |  |
| Method | AUROC | AUPRC | Precision | Recall | Specificity | MCC | Year |
| PNAbind<sup>A</sup> | **0.916** | **0.378** | **0.337** | **0.618** | **0.927** | **0.413** | 2023 |
| PNAbind<sup>M</sup> | 0.896 | 0.314 | 0.294 | 0.613 | 0.915 | 0.378 |  |
| GraphBind<sup>17</sup> | 0.854 | -- | 0.294 | 0.463 | -- | 0.322 | 2021 |
| NucleicNet<sup>35</sup> | 0.788 | -- | 0.201 | 0.371 | -- | 0.216 | 2019 |
| aaRNA<sup>34</sup> | 0.771 | -- | 0.166 | 0.484 | -- | 0.214 | 2014 |
| DNA-129 dataset |  |  |  |  |  |  |  |
| Method | AUROC | AUPRC | Precision | Recall | Specificity | MCC | Year |
| PNAbind<sup>A</sup> | **0.953** | **0.566** | **0.495** | **0.731** | 0.953 | **0.571** | 2023 |
| PNAbind<sup>M</sup> | 0.942 | 0.547 | 0.488 | 0.671 | **0.955** | 0.540 |  |
| GraphSite<sup>16</sup> | 0.934 | 0.544 | 0.460 | 0.665 | 0.950 | 0.519 | 2022 |
| GraphBind<sup>17</sup> | 0.928 | 0.519 | 0.422 | 0.684 | 0.941 | 0.500 | 2021 |
| DNA-181 dataset |  |  |  |  |  |  |  |
| Method | AUROC | AUPRC | Precision | Recall | Specificity | MCC | Year |
| PNAbind<sup>A</sup> | **0.931** | 0.333 | 0.311 | 0.632 | 0.937 | **0.409** | 2023 |
| PNAbind<sup>M</sup> | 0.921 | 0.313 | 0.293 | **0.634** | 0.932 | 0.395 |  |
| GraphSite<sup>16</sup> | 0.917 | **0.369** | **0.354** | 0.517 | **0.958** | 0.397 | 2022 |
| GraphBind<sup>17</sup> | 0.904 | 0.339 | 0.293 | 0.624 | 0.933 | 0.392 | 2021 |

### Mechanistic Interpretation of DNA versus RNA Binding Predictions

Motivated by the high accuracy of our models for distinguishing DNA binding from RNA binding, we sought to better understand how our models learn to separate these two classes of proteins. We present two methods of interpreting our model predictions, based on feature and spatial attribution, and relate these to underlying physical mechanisms that determine DNA and RNA binding.

### Feature Attribution

Feature attribution quantifies how much a particular feature or group of features determines the predictive capability of a model. In our approach, we use feature permutation^31^ whereby a group of features are permuted (along the vertex dimension of the surface mesh) such that the spatial distribution over the protein surface is mixed up. A visual example of the permutation process is shown in Fig. 4a. An error measure is computed for the permuted and unpermuted input, and the error introduced by permuting a group of features is called the feature importance, with more important features producing larger error upon permutation. Fig. 4b shows the importance of feature groups for models trained on the three datasets described above, with AUROC used to measure decrease in performance. In all cases, a drop in performance is observed, indicating that all feature groups contribute some information related to binding function. On the task of predicting DBP versus nBP, hydrogen bond donor/acceptors (HB), Atchley factors (AF), and electrostatic potential features (EP) stand out as most important. For RBP versus nBP, AF and EP show high importance, with much less importance assigned to HB. In distinguishing DBP versus RBP, AF, HB and EP features are all assigned high importance, but more strikingly geometric features (include edge features EF) show much higher relative importance, with the geometric shape descriptor HKS being assigned more importance than EP. This suggests that while electrostatic and chemical signatures may be sufficient to predict DNA and RNA binding from non-binding, the geometry of the protein surface is crucial in order to distinguish DNA from RNA binding.

**Fig. 4:**
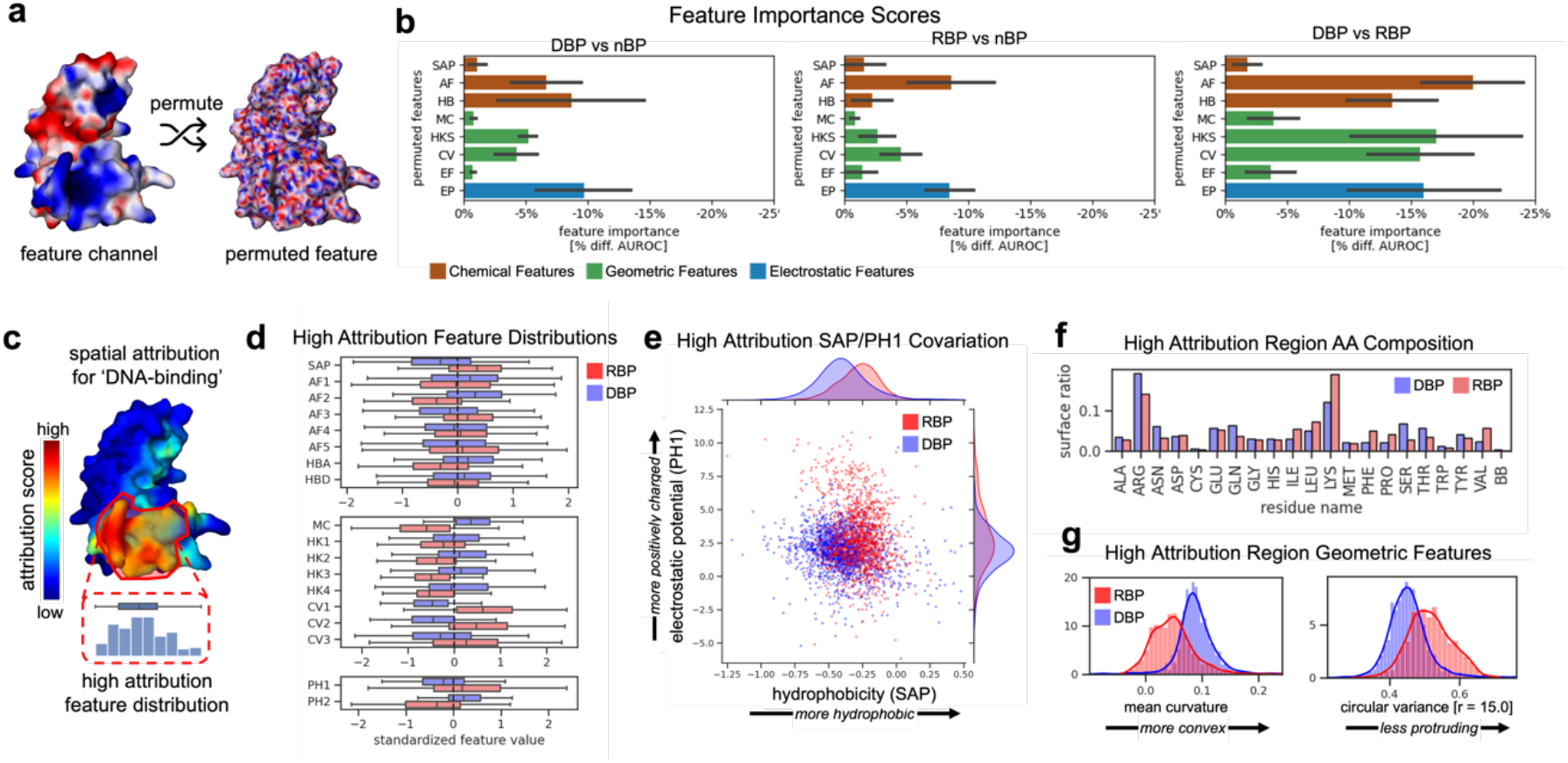
Binding function prediction results on AlphaFold2 protein structures. **a** A visual example of a permuted feature for the AlphaFold2 structure of repressor CtsR (UniProt ID C3W947). **b** Feature importance scores for the three classification models. The more negative the score the more important the feature is for contributing to model performance (as measured by AUROC). **c** A visual example of spatial attribution scores for the ‘DNA-binding’ class of the repressor CtsR (UniProt ID C3W947). **d** Feature distributions within high attribution regions for DBP and RBP proteins. **e** The co-variation of spatial aggregation propensity (SAP) and electrostatic potential (EP) in high attribution regions of DBP and RBP, showing distinct chemical and electrostatic signatures in these proteins. Every point corresponds to the mean value for a single protein. **f** Examples of geometric feature distributions showing notable differences between DBP and RBP. **g** Proportion of residue side chains and backbone (BB) within high spatial attribution regions of DBP and RBP.

### Spatial Attribution

Additional insight can be gained by performing spatial attribution, which quantifies how strongly different regions of the protein molecular surface contribute to the predicted probability of a given binding class. Intuitively, regions of a protein structure that most affect model predictions should be related to the binding mechanism of the protein, and correlate with the nucleic acid binding site. We computed the spatial attribution via Grad-CAM^32^, an attribution method where gradients for a target class probability are computed with respect to the activations of a chosen convolutional layer (typically the final layer), and average-pooled along the feature map dimension to produce a localized score for each element of the activation. These scores quantify how strongly a region of the input affects the predicted probability of a target class.

Regions of low attribution contribute little to the predicted probability of a given class, and regions of high attribution strongly determine the output probability. In Fig. 4c we show a visual example of the attribution scores for the ‘DNA-binding’ class from the DBP versus nBP model for the repressor CtsR (UniProt ID C3W947). The attribution scores for the same protein are shown in Supplementary Fig. 2a alongside the known DNA binding site (determined from a native co-crystal structure), showing that regions of high attribution are visually correlated with the DNA binding site. In Supplementary Fig. 2b. we show that in general, regions of high spatial attribution for either NA binding class across all models correspond to regions that lie predominantly within known NA binding sites. DBP and RBP in our AlphaFold2 datasets which had experimentally determined co-crystal structures available in the PDB were identified and the binding sites of these proteins were labeled using the co-crystal. We then computed precision and recall curves based on the attribution scores (first normalized between zero and one).

These curves show that while the high attribution regions have low recall, they attain good precision – meaning the high attribution regions reside predominantly within the observed binding sites. This also indicates only particular regions within the binding sites are necessary for binding prediction.

Next, we plotted the distribution of input features within high attribution regions for every test set protein, using attributions scores computed from each of the three classification models described above. The threshold for high attribution was arbitrarily set at the 75^th^ percentile across all values for a given protein. Scores were computed for the class corresponding to the ground-truth label of each protein. High attribution feature distributions are shown in Fig. 4d for the DBP versus RBP model, and for all models and features in Supplementary Fig. 2c-d. Clear separation can be seen in the distributions of many features for each target class and model.

We note that in almost every case, feature distributions with large separation between classes also correspond to feature groups which were assigned a high importance as shown in Fig. 4b. The high attribution feature distributions provide insight into how different features may be related to binding mechanisms. For the DBP versus RBP model, clear separation is seen in the features SAP (hydrophobicity) and PH1 (electrostatic potential). A scatter plot of the covariation of these features in Fig. 4e shows that the high attribution regions of RBP surfaces tend to be both less hydrophilic but more electrostatically positive compared to DBP proteins. Lysine and arginine occur frequently in NA binding sites as both are positively charged and form favorable electrostatic interactions with the negatively charged phosphate groups in the NA backbone.

However, the proportions of these side chains may differ as we observe in Fig. 4f. The high attribution regions of RBP show a higher lysine content, while those of DBP show a higher arginine content. Lysine is less hydrophilic than arginine, explaining the difference in hydrophobicity observed. We note that this observation is consistent with a meta-analysis of DNA and RNA binding residues performed by Zhang et al.^33^ indicating a marginal preference for lysine in RNA binding sites and a preference for arginine in DNA binding sites (Table 3 of reference). The enhanced electrostatic potential seen for RBP may be necessary for stabilizing the more globular structure RNA tends to adopt in protein-RNA complexes (see Fig. 5a for an example), which would otherwise experience destabilizing electrostatic repulsion between phosphate groups of the RNA backbone.

**Table 3:**
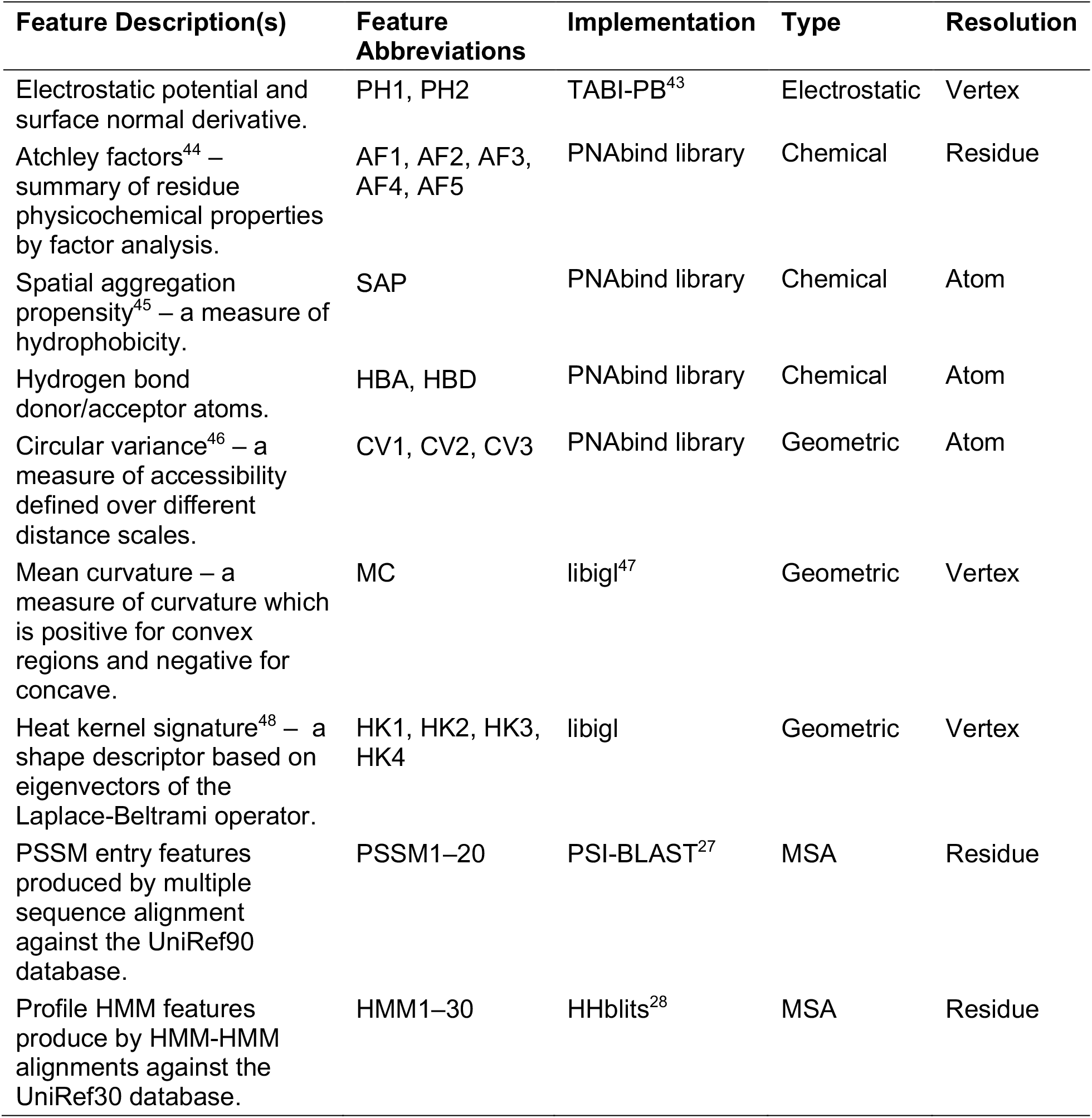
A complete list of vertex features used in our method. Feature abbreviations are shorthand for every feature name used in figures throughout the manuscript. Implementation is any external packages or software which PNAbind uses to implement the respective feature. The references under this column refer to the reference where the implementation was described. Type refers to which group a feature belongs to – electrostatic, chemical, geometric or multiple-sequence alignment (MSA). Resolution refers to what structural resolution the feature is naturally defined for, which can be ‘Residue’, ‘Atom’, or ‘Vertex’.

| Feature Description(s) | Feature Abbreviations | Implementation | Type | Resolution |
| --- | --- | --- | --- | --- |
| Electrostatic potential and surface normal derivative. | PH1, PH2 | TABI-PB<sup>43</sup> | Electrostatic | Vertex |
| Atchley factors<sup>44</sup> – summary of residue physicochemical properties by factor analysis. | AF1, AF2, AF3, AF4, AF5 | PNAbind library | Chemical | Residue |
| Spatial aggregation propensity<sup>45</sup> – a measure of hydrophobicity. | SAP | PNAbind library | Chemical | Atom |
| Hydrogen bond donor/acceptor atoms. | HBA, HBD | PNAbind library | Chemical | Atom |
| Circular variance<sup>46</sup> – a measure of accessibility defined over different distance scales. | CV1, CV2, CV3 | PNAbind library | Geometric | Atom |
| Mean curvature – a measure of curvature which is positive for convex regions and negative for concave. | MC | libigl<sup>47</sup> | Geometric | Vertex |
| Heat kernel signature<sup>48</sup> – a shape descriptor based on eigenvectors of the Laplace-Beltrami operator. | HK1, HK2, HK3, HK4 | libigl | Geometric | Vertex |
| PSSM entry features produced by multiple sequence alignment against the UniRef90 database. | PSSM1–20 | PSI-BLAST<sup>27</sup> | MSA | Residue |
| Profile HMM features produce by HMM-HMM alignments against the UniRef30 database. | HMM1–30 | HHblits<sup>28</sup> | MSA | Residue |

**Fig. 5:**
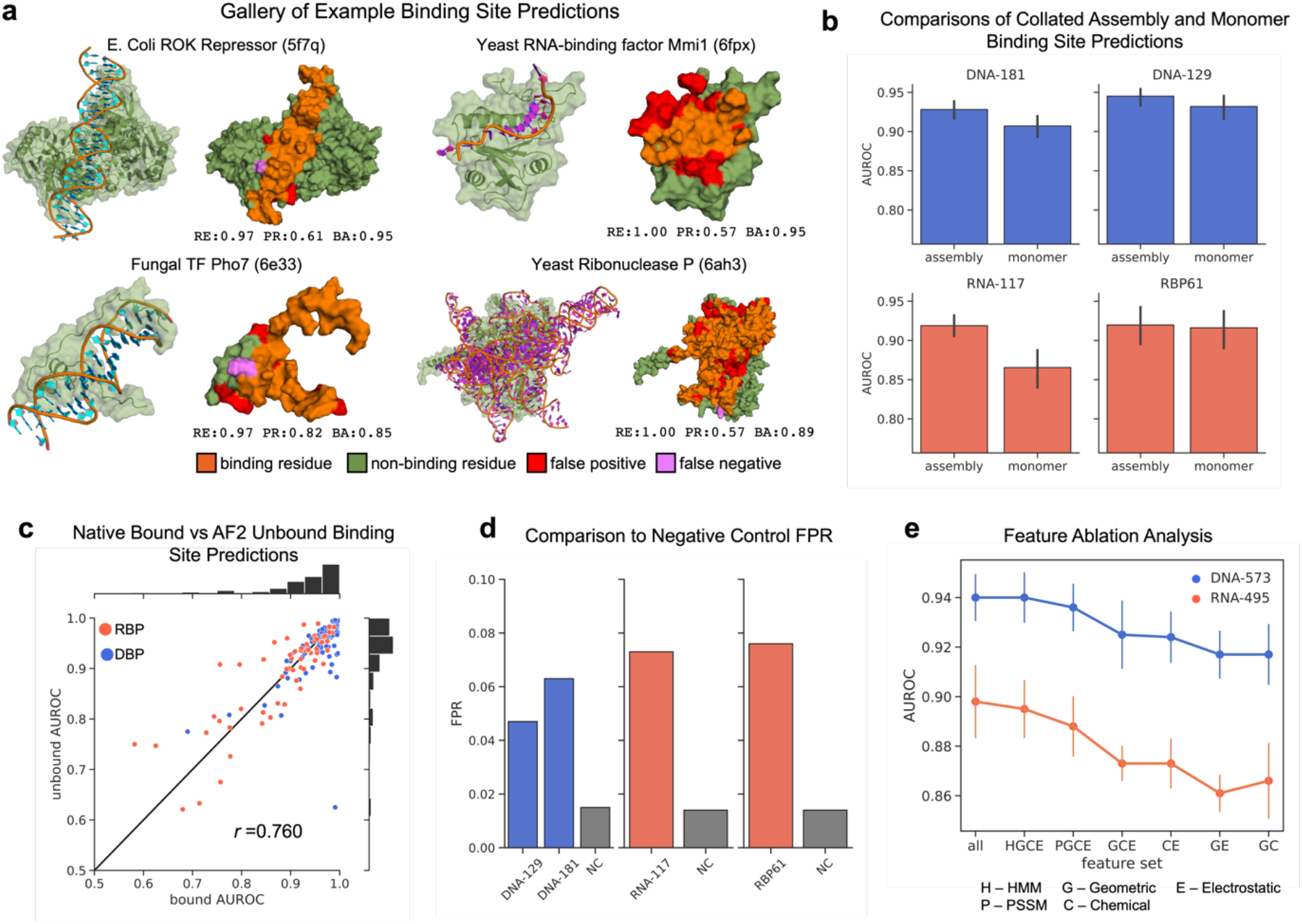
Binding site prediction results on native protein structures. **a** Gallery of example binding site predictions from test sets. Select DNA (RNA) binding proteins are shown bound to target DNA (RNA) alongside their binding site predictions. Corresponding performance metrics are given – recall (RE), precision (PR) and balanced accuracy (BA). **b** A comparison of model accuracy for binding site predictions made on target chains in their native structural context given by the full biological assembly versus predictions made on single-chain structures with no additional structural context. Only collated surface residues were used in these comparisons. **c** AUROC of binding site predictions for bound native structures versus AUROC of binding site predictions for in silico unbound structures across proteins from the four test sets. The solid black line indicates perfect agreement. **d** Comparison of the overall false positive rates of the three binding site prediction models on a negative control (NC) dataset and benchmark test sets. **e** Cross-validated AUROC scores for various feature sets showing how the model accuracy decreases as certain features are removed.

Additionally, differences in geometric feature are notable, as shown in two geometric feature distributions in Fig. 4g. These features are both measures of curvature defined over different distance scales. The DBP distribution shows a shift towards more positive mean curvature, which indicates that high attribution is assigned to regions with more convexity compared to RBP. The DBP distribution also shows a shift towards lower values of CV2 (circular variance with a radius of 15.0; see Methods). A low value of this feature indicates a point on the surface is protruding outward relative to other regions of the surface within a chosen radius. Taken together, these distributions are consistent with protrusions from the molecular surface on the order of the width of the DNA major and minor grooves. We hypothesize that the differences in geometric features of DBP versus RBP proteins within high attribution regions is related to the fact that DNA predominantly adopts a B-form double helix, while RNA folds into much more globular structures or adopts an A-form double helix, and these differences in the tertiary structure of DNA and RNA are reflected in the geometry of the protein molecular surface.

### Binding Site Prediction

We trained models for residue-level binding site prediction using our graph-segmentation network and validated the accuracy of these models on previously developed benchmark datasets. Three independent sets of models were trained (using identical hyperparameters; see Fig. 2 and Methods) on DNA binding protein and RNA binding protein training sets (DNA-573^17^, RNA-495^17^, RBP09^24^ – Supplementary Fig. 4a). Training was performed using fivefold cross-validation, and an ensemble consisting of the best model from each training fold was used for predictions. Trained models were then validated using four test sets, two containing native DNA binding proteins, and two native RNA binding proteins (DNA-181^16^, DNA-129^17^, RNA-117^17^ and RBP61^24^ respectively – Supplementary Fig. 4a). Each test set was designed previously to have minimal sequence overlap with a corresponding training set or to only include co-crystal structures determined more recently than any structure in a corresponding training set.

### Performance and Comparison with Recent Methods

Validation metrics for our method and other recently developed methods are given in Table 2. We evaluated our models using the same residue label criteria used previously in the cited studies. Namely, DNA-129, RNA-117, and DNA-181 binding site labels were constructed using interaction annotations from the BioLiP database^16,17^ and RBP61 residue labels were constructed based on a nucleotide-residue distance threshold of 5.0 Å^24^. PNAbind achieves the highest AUROC for all four test sets, ranging from 0.916-0.953. Two rows are shown for PNAbind, the first one indicates the performance of our models on the biological assembly of the protein, and the second on the monomeric state (e.g., single-chain, irrespective of the native binding mode of the protein). Fig. 5a shows a gallery of binding site predictions on examples drawn from the test sets, with corresponding metric values for those predictions. Our models accurately identify DNA and RNA binding sites for a wide variety of structural domains and can detect binding sites of proteins with specificity for single-stranded, double-stranded and more complicated secondary structures. The proteins used for validation have a diverse range of biological functions, as shown in Supplementary Fig. 4c.

### Predictions Made Using Protein Assemblies Are More Accurate

In Table 2, two rows are given for PNAbind. The first shows validation metrics for models evaluated on the native experimental biological assembly of the protein, and the second on the monomeric structure (e.g., a single protein chain, irrespective of the native binding mode of the protein). On the benchmark datasets the full-assembly predictions achieve consistently higher validation metrics than the monomeric predictions. To assess if the native assembly provides an inherent advantage over the monomeric structure for binding site prediction, we collated all surface residues common to both structural models. The collation ignores residues from the monomeric structure that participate in protein-protein interactions and become buried beneath the solvent-accessible surface in the assembly structure. Therefore, the collated residue sets provide a direct one-to-one comparison. The validation metrics on these collated datasets are shown in Fig. 5b. In agreement with the values in Table 2, the assembly structure consistently has better AUROC than the monomeric structure. This indicates that PNAbind can capture not only the local chemical, evolutionary and geometrical properties of the protein, but also the structural context in which a binding site might appear in relation to co-factors, dimerization etc. This suggests that where available, the full native assembly should be used for binding site predictions.

### Binding Site Predictions on AlphaFold Predicted Protein Structures

The binding site prediction models described above were trained and validated on native protein structures bound to their nucleic acid targets. However, the conformation of a protein may change upon binding, and we wanted to address the question of how much conformational variation between the bound and unbound state of a protein may affect the accuracy of our model predictions. Many NABP lack native structures in the unbound state, so we used AlphaFold2 predicted structures as a proxy for native unbound structures. DNA and RNA binding residues in the unbound proteins were labeled via mapping binding site labels from the bound native structure. The mapped binding site labels allowed us to estimate the agreement of predicted binding sites between bound and unbound structures. Our results are shown in Fig. 5c, where we plot the AUROC for every pair of bound/unbound structures. Good agreement in AUROC is seen for both native and AlphaFold2 derived structures, with an overall Pearson correlation of 0.76. We note that in general the sequence alignment between native and predicted structure is not perfect because the predicted structure corresponds to the wild-type sequence, and the sequence used in the experimental model may contain modifications in order to overcome solubility or crystallization difficulties, and we expect that these sequence modifications contribute in part to some of the discrepancies observed. Overall, however, the high correlation demonstrates that our method is applicable to both bound and unbound protein structures.

### Negative Control Experiment

The training and testing datasets used for binding site prediction sample a wide variety of proteins with diverse structure and functional roles related to NA binding (Supplementary Fig. 4c). However, because the non-binding site regions are sampled only from structures of NABPs, it is not clear how well the full domain of the non-binding feature space (e.g., the full space of protein structures which have no NA binding function) is sampled, and how the models will behave on this domain. To measure the performance of our models on out of domain targets, we constructed a negative control dataset which consists of proteins with no known NA binding function (see Methods). For these proteins, we consider any binding site prediction to be a false positive, and we assess the model performance on the negative control using false positive rate (FPR).

Fig. 5d shows the FPR rates on the negative control dataset compared the FPR on the benchmark test sets. In each case, the FPR of the negative control is significantly lower than that observed for test set NA binding proteins. This indicates that our models perform better than expected on proteins which are outside the training domain (in a functional sense). We hypothesize that this is largely because our models capture physical mechanisms of protein-NA binding, providing a high degree of generalizability. Supplementary Fig. 3 shows the FPR of each model for each residue type on the negative control and test sets. Variation is seen across the models and datasets, but in general, positively charged side chains (arginine, lysine, histidine) and aromatic side chains (tyrosine, tryptophan, phenylalanine) have the highest FPR.

### Feature Ablation Study

We performed a feature ablation study to measure the contribution of different groups of features to the accuracy of our models. Fig. 5e shows fivefold cross-validated AUROC scores for two of the benchmark training datasets. The labels and counts of the feature groups are PSSM features (P, 20), HMM profile features (H, 30), chemical features (C, 8), geometrical features (G, 8) and electrostatic features (E, 2). The highest AUROC scores are achieved when all groups of features are included. Among the MSA features (P/H), the profile HMMs appear to be more informative for both DNA and RNA binding sites, but marginal performance improvement is achieved by including PSSM features.

Among the three subsets of structural features, CE features appear more informative of discriminating DNA/RNA binding sites from non-binding sites, consistent with the findings in Fig. 4b for models trained to predict binding function.

### Case Study – APOBEC3G Dimerization and RNA Binding

APOBEC3G (A3G) is a cytidine deaminase that catalyzes the conversion of cytosine to uracil on DNA and RNA substrates via a conserved zinc-coordinating motif that forms a catalytically active binding pocket. It is a member of the APOBEC3 (A3) family of proteins which play important roles in mammalian innate immune response against retroelements and retroviruses including HIV-1^36-38^. A3G has a strong affinity to bind RNA in multiple binding modes^37^ and contains two domains known as CD1 and CD2. Yang et al.^37^ published the first structures of full-length A3G from monkey rhesus macaque (*Macaca mulatt*) in its homodimer configuration that is mediated through CD1-CD1 interactions. Although the protein was co-purified with bound RNA, they were unable to construct a structural RNA model based on the observed electron density. Using these recently determined A3G structures, we applied our models (trained on the RNA-495 dataset) to predict RNA binding in both the monomer and dimer conformation of A3G to determine if our models could help elucidate the role RNA binding plays in the dimerization of A3G.

Fig. 6c demonstrates RNA binding site predictions on full-length A3G (CD1+CD2) in the dimer and monomer configuration. Yang et al. solved three structures of the A3G homodimer at different pH (PDB IDs: 6P3X, 6P3Y, 6P3Z). The predictions shown are averaged over residue labels predicted from all three structures. Our models predict a large RNA binding region (marked region 1) that spans the CD1-CD1 dimerization interface in the dimer structure, which is not present in the monomer structure. This is consistent with the experimental observation that A3G dimerization is RNA-dependent^37^, and agrees with the region Yang et al. hypothesized may form the dimerized RNA binding site (Yang et al., Fig. 2). Yang et al. mutated critical residues in the dimerization interface of A3G which were shown to disrupt the dimerization and reported their experimental data (shown in adapted form in Fig. 6b). Their data demonstrates a dramatic decrease in RNA association for the constructs with dimerization-disrupting mutations (rM9, rM10, rM15). Our models also predict a second, smaller region of RNA binding (marked region 2) that is independent of dimerization (e.g., is predicted for both monomer and dimer structures) and suggests that A3G should still possess some RNA binding capacity even if dimer formation is disrupted. This is consistent with the results reported by Yang et al. who measured binding affinities for 50-nt ssRNA for all mutants listed in Fig. 6b.

**Fig. 6:**
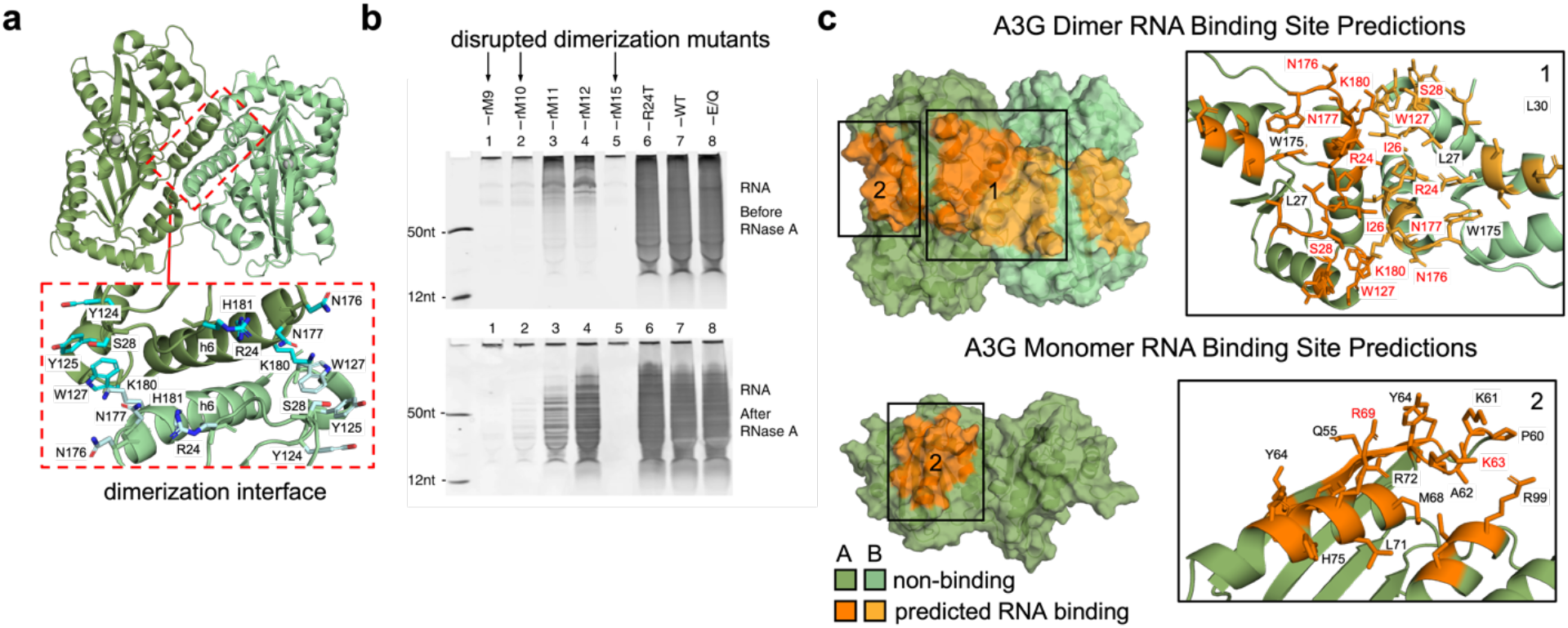
RNA binding site predictions for the APOBEC3G protein. **a** The structure of full-length A3G from monkey rhesus macaque (*Macaca mulatt*) is shown as a homodimer (PDB ID 6P3Y). The dimerization interface is shown with residues from a-helix h6 and adjacent loops that participate in dimerization (adapted from Yang et al.^37^). **b** Denaturing urea polyacrylamide gel analysis (adapted from Yang et al.^37^, Fig. 3) of RNA molecules associated with WT and mutant A3G proteins both before and after RNase A treatment. Mutants which have disrupted dimerization are indicated. Reduced RNA association for these constructs is observed in the gels. All data collected was by Yang et al. **c** RNA binding site predictions for A3G. The top panel shows predicted binding regions in the dimer conformation and the bottom panel shows predictions in the monomer conformation. Our models predict a large RNA binding region (marked as region 1) that spans the CD1-CD1 dimerization interface which is not present in the monomer. A smaller RNA binding region (marked as region 2) is predicted for both conformations. The insets for region 1 and 2 indicate residue names in each region. Residues names shown in red are those which Yang et al.^37^ identified as important for RNA binding.

Yang et al. performed mutational studies to identify residues which are relevant for RNA binding. They determined several residues that play a key role in RNA binding which overlap with our model predictions. In region 1, these are R24, I26, S28, W127, N176, N177, and K180, and in region 2, K63 and R69. These residues are indicated in the insets for regions 1 and 2 in Fig. 6c. The results of Yang et al. provide strong experimental corroboration of our model predictions for APOBEC3G RNA binding. Overall, our results suggest that RNA binding to region 1 and dimerization of A3G may be mutually dependent on each other, and loss of dimerization may consequently result in loss of RNA binding to the resulting half-site of region 1.

## Discussion

PNAbind predicts global protein binding function (e.g., DNA-binding versus RNA-binding) or predicts individual binding residues for binding site prediction, using surface-based representations of proteins. The corresponding choice of features is motivated by aspects of the protein molecular surface geometry and chemistry related to known NA recognition mechanisms. The geometry of the protein molecular surface is an important feature to consider and has been underutilized in previous works. Structural motifs found within the binding interface may have geometries related to binding function, such as binding pockets that can capture nucleotide bases in ssDNA, or binding elements suitable for insertion into the major or minor groove of dsDNA. Our results on discriminating DNA versus RNA binding function highlight the importance of geometric information, as our models show geometric features to be at least as important if not more so than the chemical and electrostatic features used.

Experimentally determined structures are currently required for training models to predict NA binding sites with high accuracy, as there are no reliable alternatives for labelling the binding site locations in a training set. Functional annotations, however, do not require any structure data and are much more abundant. It is noteworthy that our models trained on predicted protein structures (many of which have no experimental structural models) achieve both high accuracy in identifying NA binding and discriminating DNA from RNA binding, but also can partially identify the binding sites of these proteins as shown in Supplementary Fig. 2a. This is achieved simply by inspecting which regions of the protein structure most contributed to the model prediction, using well-established attribution methods^32^. This approach allows the models to be applied to individual proteins for better understanding of how the protein structure relates to the binding function of the protein, or to perform a large-scale analysis and compare differences between classes of proteins as we have done in this study. A major strength of our approach is that it achieves high accuracy of its predictions but also allows for interpretation of model predictions in terms of structural features and corresponding binding mechanisms.

A significant majority of known protein sequences are poorly annotated. With over 200 million predicted protein structures now available in the AlphaFold database^29^, PNAbind can be used to aid in high-throughput annotation of NABPs. Our binding site prediction models provide high accuracy on a more specialized task and can be used to aid in interpretating biochemical data, identify functionally important residues, or provide prior information about binding sites for modeling of protein-NA complexes. We also note that, while we have focused exclusively on nucleic acid binding in this report, PNAbind is a structure-based approach and hence is quite general and can be applied to the prediction of other types of functional annotations or other protein-ligand binding sites. Our method provides a general way to characterize how structural properties of proteins are related to their biological function, and in principle can be applied to any class of proteins.

## Methods

### Mesh Generation

We use NanoShaper^39^ for generating the molecular surface mesh and found the ‘skin’ surface^40^ worked well in our study. The smoothness of the skin surface is controlled by a scalar ‘shrink factor’, which we set to 0.45, producing a surface that is smooth but captures geometrical features such as binding pockets. We use van der Waals parameters from the AMBER ff99 force field^41^. Prior to mesh generation, missing atoms are added to the structure using PDB2PQR^42^. The AMBER99 parameter set defines a radius of 0 for some hydrogen atoms – in these cases we set the radius to a minimum value of 0.6 Å.

### Vertex Features

A list of vertex features used in our study is given in Table 3. Different features are computed at different resolutions before being mapped to the surface mesh. For example, MSA features are computed at the level of individual residues, hydrogen bond donor/acceptor labels are assigned to individual atoms, and the surface mean curvature is computed for mesh vertices. Residue-level features are first mapped to child atoms, and atom-level features are mapped to vertices using a distance weighted average, with weights given by the inverse Euclidean distance between the atom and each nearby vertex, for a maximum distance cutoff of 2.5 Å.

PH1 and PH2 refer to the electrostatic potential and the normal component of the electric field, respectively. The electrostatic potential is computed using TABI-PB^43^, a boundary integral Poisson-Boltzmann solver. We set the bulk ion concentration to 0.15 mol and the interior dielectric constant to 2.0. AF1–5 are numerical descriptors known as Atchley factors that describe a variety of physicochemical attributes such as polarity, secondary structure propensity, molecular volume, codon diversity, and electrostatic charge of amino acids. SAP is short for spatial aggregation propensity and measures hydrophobicity. HBA and HBD indicate proximity to a hydrogen bond acceptor or donor. CV1–3 is a geometric feature known as circular variance that roughly corresponds to the solid angle of a sphere (centered on each vertex) that is occupied by the molecular volume of the protein. For points which protrude from the surface, most of the sphere is unoccupied and the feature takes a value near zero. For points, which are buried deep within a pocket or channel, almost all the volume of the sphere is occupied, and the features takes a value near 1. Each CV feature is defined with respect to a sphere of radius 7.5,

15.0 and 30.0 Å respectively. MC stands for mean curvature, a standard geometric measure of curvature that is positive for convex surfaces and negative for concave surfaces. HK1–4 are heat kernel signatures (HKS), widely used shape descriptors defined in terms of eigenvectors and eigenvalues of the Laplace-Beltrami operator. They are mathematically related to the diffusion of heat over a surface, and in practice are good descriptors of shape that are robust to small perturbations to the isometry of a surface. Each HKS feature encodes geometric information over difference distance scales as shown in Supplementary Fig. 1. PSSM1–20 are features derived from PSSM matrices computed by PSI-BLAST. Sequences of target chains are searched against the UniRef90 database with an E-limit of 10, E-value cutoff of 0.001, gap open penalty of 1, gap extend penalty of 1 and 3 iterations. HMM1–30 are features derived from profile HMMs computed by HHblits. Sequences of target chains were searched against the UniRef30 database using default run parameters.

### Edge Features

We use the same edge features described by Deng et al.^49^, which include the edge length (vertex-vertex distance), the angle an edge makes with the surface normal at each adjacent vertex, and the angle between the surface normal at adjacent vertices. These features encode rotationally invariant geometric information about how the vertices within a radial neighborhood are distributed in space. For example, if all vertices lie in a plane, the normal-normal angles will be zero, and all edge-normal angles will be 90*°*. However, if the vertices lie on a sphere, then the normal-normal and edge-normal angles will vary as a function of distance in a systematic way.

### Probabilistic Binding Model

Given a mesh representation of a protein molecular surface, *G* = {**V** ∈ ℝ^|*V*|×3^, **E** ∈ ℕ^|E|×2^} where **V** are the vertex coordinates in Euclidean space, **E** are pairs of indices into **V** that form directed edges corresponding to triangle faces, **X**_*V*_ are vertex features and **X**_E_ are edge features, we wish to either classify the entire graph based on the global aggregation of local physicochemical properties, or to classify every vertex *ν*_*i*_ ∈ **V** as a binding site or non–binding site based on features of that vertex and features of its local environment. Both approaches share similar frameworks, and we first describe the framework for vertex classification (i.e., segmentation).

### Segmentation

We compute a per-vertex Bernoulli distribution *q*(*y*_*j*_|*G*, **X**_*V*_, **X**_E_; **θ**) ∈ [0,1] where *y*_*i*_ is the class label of *ν*_*i*_; *y*_*j*_ = 1 represents a binding site and *y*_*i*_ = 0 a non–binding site. The distribution *q* depends on a parameterization **θ** defined by the neural network. We note that **θ** does not depend on *i* — therefore the parameters are shared over all vertices in *G*.

Let *𝒟* = {*G*_*n*_, ***p***_*n*_} be a dataset of protein molecular surface graphs and ***p***_*n*_ are the corresponding per-vertex binding probabilities, *p*_*ni*_ ∈ {0,1}. *p*_*ni*_ = 1 if the *i*^th^ vertex lies in an observed nucleic acid binding site and is 0 otherwise. We learn *q* by minimizing the cross-entropy of our predicted probabilities ***q*** with the observed probabilities ***p***_*n*_ over all graph vertices. That is, we seek to find a set of parameters that satisfies the following condition,

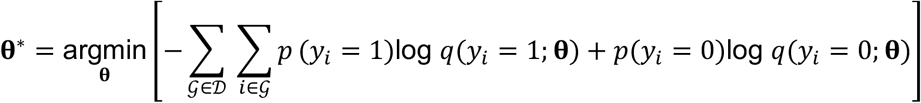

Given an estimate for **θ**^*\**^, we can predict the probability for every vertex in *G* to belong to a binding site or not, and assign labels to each vertex by choosing a threshold on *q*_*i*_,

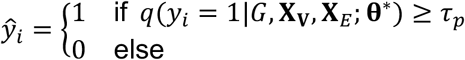

where *ŷ*_*i*_ is the predicted label for the *i*^th^ vertex and *τ*_*p*_ ∈ [0,1] is a threshold value that can be chosen depending on the application. In practice **θ**^*\**^ is estimated via gradient descent.

### Graph Classification

To assign a probability to the entire graph, we modify our learned distribution to be of the form *q* = *q*(*y*|*G*, **X**_*V*_, **X**_E_; **θ**) where *y* is now a single binary class label for the entire graph, and our dataset is of the form *𝒟* = {*G*_*n*_, *p*_*n*_} where *p*_*n*_ ∈ {0,1}. The model parameters can be optimized in the same way described above by minimizing a cross-entropy loss function using gradient descent.

### Probability Thresholds

Assigning a class given a probability depends on choosing a threshold which is in practice somewhat arbitrary. For segmentation, due to the imbalanced nature of the datasets (statistics are shown in Supplementary Fig. 4a), an unbiased prior of 0.5 may be a sub-optimal choice. We applied Platt scaling to each model (using validation folds for scaling parameter optimization) before combining probabilities in the ensemble and chose a probability threshold that maximized the F1 score of training set predictions, such that each ensemble model has an independent threshold assigned to it. We did not apply Platt scaling to binding function prediction models and chose a threshold of 0.5 for those models, as these datasets are well balanced.

Segmentation models produce vertex-level binding site probabilities. Residue-level probabilities are obtained by max pooling over all vertices that correspond to the solvent-excluded surface of each residue.

### GNN Model Architecture Segmentation

We utilize convolutional graph neural networks (GNN) to model per-vertex binding site probabilities 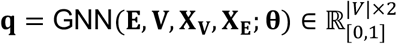. The network is based on a graph U-Net^25^ style architecture, combined with the iterative farthest point sampling (FPS) pooling and *k*-nearest neighbors (KNN) unpooling used in PointNet++ by Qi et al.^26^ The crystal-graph (CG) convolution introduced by Xie et al.^50^ was chosen for the convolution kernel for all convolutional layers. The network architecture is shown in Fig. 2b. Fig. 2c shows a schematic overview of how convolutional, pooling and unpooling layers work. First a uniformly distributed subset of vertices is sampled via FPS, with a sampling ratio of *α* = 0.5. Centered on each sampled vertex (referred to as a centroid) a directed radial graph is constructed by joining each centroid to all other vertices within a sphere of radius *r*. A message passing convolution operation is then performed over this radial graph. The incoming messages for every centroid are aggregated via a linear combination of min, max, mean and stdev pooling and the final activation for each centroid is given by the sum of the pooled messages and the current centroid activation. Updated centroid activations contain information about the spatial context of each centroid within the neighborhood defined by the radius *r*. Convolutions are repeated twice per layer, after which non-centroid vertices are removed from the graph and the centroids form a new, sparser set of vertices. The sampling/convolution/pooling operation is then repeated *d* = 3 times. We choose increasing radii for each convolutional layer in the network with *r* = 5.0, 7.5, 10.0 Å. To perform segmentation pooled vertices must be interpolated back to restore the original resolution of the mesh, which is accomplished by KNN unpooling^26^. Starting from the final coarsened layer ℓ = ℓ_6_, we interpolate features of pooled non–centroid vertices 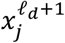 from the *k* = 3 nearest centroid vertices as shown in Fig. 2d. The interpolated features are a distance-weighted mean of centroid features. Finally, these interpolated features are combined with the matching vertex activations from the corresponding convolution/pooling layer via skip-connections as shown in Fig. 2b. The skip connections improve gradient flow through the network and allow spatial context information from earlier layers to propagate forward more efficiently. The output of the final unpooling layer is concatenated with the full-resolution skip connections and passed through a two-layer MLP which acts like a 1×1 convolution to transform the activations to ℝ^2^. After softmax normalization, the output of the model represents a Bernoulli distribution over each vertex.

### Classification

The architecture of the classification model is identical to the segmentation model up to the final convolutional/pooling layer and is shown in Fig. 2a. In place of unpooling layers, a single global pooling layer is used to aggregate all information over the coarsened mesh to a fixed-dimensional activation. The global attention pooling is of the form

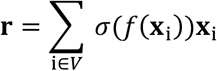

Where *f*(·) is an MLP with learnable weights and *σ*(·) is the sigmoid function.

### Datasets Classification

We developed three datasets for training models to predict DNA and RNA binding function using structures based on AlphaFold2 predictions. Three sets of proteins were identified in the Swiss-Prot^51^ knowledgebase using functional annotations – DNA binding proteins (DBP), RNA binding proteins (RBP) and non-binding proteins (nBP). DNA/RNA binding proteins were first identified by searching the database for sequence entries with the UniProtKB molecular function keyword ‘DNA-binding’ or ‘RNA-binding’. Any protein sequence which contained both annotations was excluded, and sequences longer than 1000 amino acids or with an annotation score less than 2 were excluded. To identify nBP, sequence entries which did not contain the keyword ‘DNA-binding’ or ‘RNA-binding’ were first identified. Entries were then excluded which contained additional keywords related to nucleic acid binding such as ‘Nucleotide-binding’, ‘Repressor’, ‘Activator’, ‘Ribonucleoprotein’ etc. Additionally, Gene Ontology (GO)^52^ molecular function annotations were also used for excluding entries that contained annotations related to nucleic acid binding. A list of more than 4400 GO terms was generated by traversing the GO annotation hierarchy starting from source terms such as ‘nucleotide binding’, ‘nucleic acid binding’, ‘nucleotide metabolic process’, ‘nucleic acid metabolic process’, etc. After filtering, AlphaFold2 derived structures were then obtained for the remaining sequences and a second round of filtering was performed based on the predicted structure quality. Low confidence regions of the predicted structures were removed (confidence < 0.65), and only structures which remained as a non-disjoint structure were kept. Finally, sequence clustering was performed on remaining sequences at a threshold of 35% sequence similarity, and two sequences per cluster were sampled. The result was three sets of proteins which we could have high confidence in both the predicted structure quality and the accuracy of the functiona annotations. These were then used for the DnP-6784, RnP-6046 and RDP-6046 datasets. Supplementary Fig. 4 shows statistics regarding these datasets.

### Segmentation

Seven benchmark structural datasets of protein–nucleic acids complexes were used in this study for training/testing segmentation models. One DBP dataset (DNA-573^17^) and two RBP datasets (RBP09^24^, RNA-495^17^) were used as training sets, and two DBP datasets (DNA-181^16^, DNA-129^17^) and two RBP datasets (RBP61^24^, RNA-117^17^) were used as independent test sets. These datasets contain experimentally derived structural models of PNA complexes deposited in the PDB^53^. Models trained on RBP09 were validated on RBP61, models trained on DNA-573 were validated on DNA-129 and DNA-181, and models trained on RNA-495 were validated on RNA-117. When using the biological assembly as the basic structural unit, any additional chains beside the target chain were masked out during training/testing for fair comparisons with previous studies. Supplementary Fig. 4 shows statistics regarding the class distribution and functional annotations of the proteins in these datasets. A negative control (NC) dataset was produced by taking the subset of the nBP dataset proteins described above which contained experimentally determined structures.

### Model Training Procedure

Model parameters were determined using the ADAM^54^ optimizer to minimize the cross-entropy between predicted probabilities and the ground-truth labels over the mesh vertices (segmentation) or entire mesh (classification). Fivefold cross-validation with early stopping was performed. For each fold, several duplicate models were trained with random initialization and within each fold the best model was selected.

## Data Availability

Data, source code and documentation for implementing and training PNAbind models is available online at https://github.com/jaredsagendorf/pnabind.

## Author Contributions

J.M.S. designed the PNAbind method conceptually, designed the GNN architecture, features, and developed code for implementing, training and evaluating models, and analyzed data. J.M.S. wrote the manuscript with help from R.R. and comments from all authors. R.M. developed code, assisted with figure preparation, implemented and tested model components, and processed data. J.H. assisted in data processing and hyperparameter tuning. X.S.C. provided data for experimental validation using the A3G protein. R.R. supervised the project.

## Acknowledgements

This work was supported by the National Institutes of Health (grant R35GM130376 to R.R.) and an Andrew J. Viterbi Fellowship in Computational Biology and Bioinformatics (to R.M.).

## Competing Interests

The authors have declared that they have no conflict of interest.

## Notes

### Competing Interest Statement

The authors have declared no competing interest.

