## Supplementary Data: Supplementary Fig. 1-4 for "PNAbind: Structure-based prediction of protein-nucleic acid binding using graph neural networks"


### Supplementary Figures and Tables

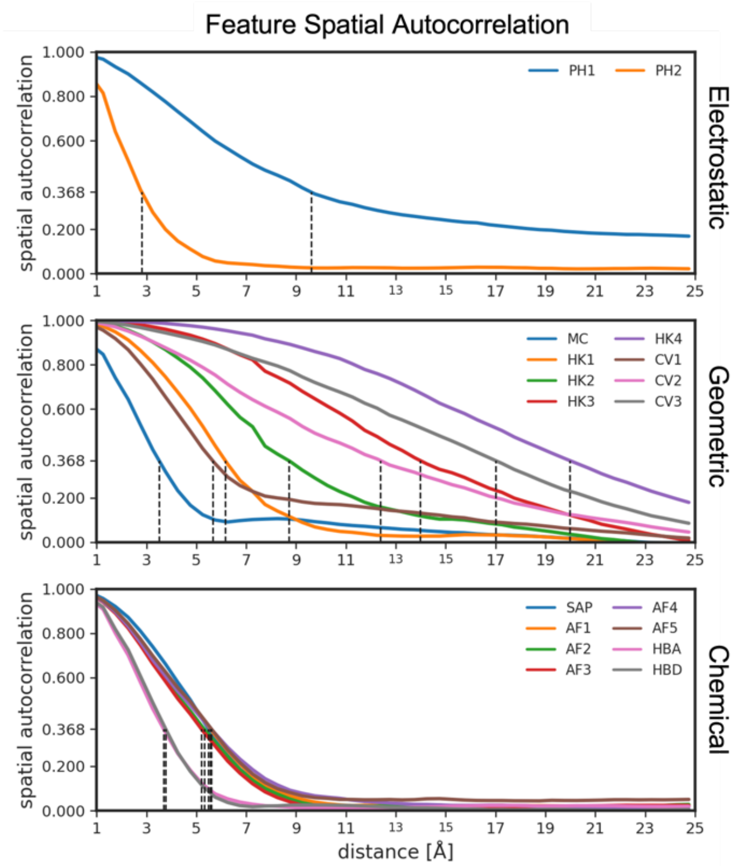


**Supplementary Figure 1: Spatial autocorrelations for vertex features.** Autocorrelations of eighteen structure-based features were computed using the Pearson correlation of feature channels between pairs of vertices within bins of increasing radial distance. Features which vary slowly over the mesh will have high autocorrelation over large distances, and features that vary rapidly have correlations that decay quickly. The dotted lines indicate the distance at which each feature autocorrelation drops to a value of $1/e$, corresponding to a characteristic distance scale for that feature.

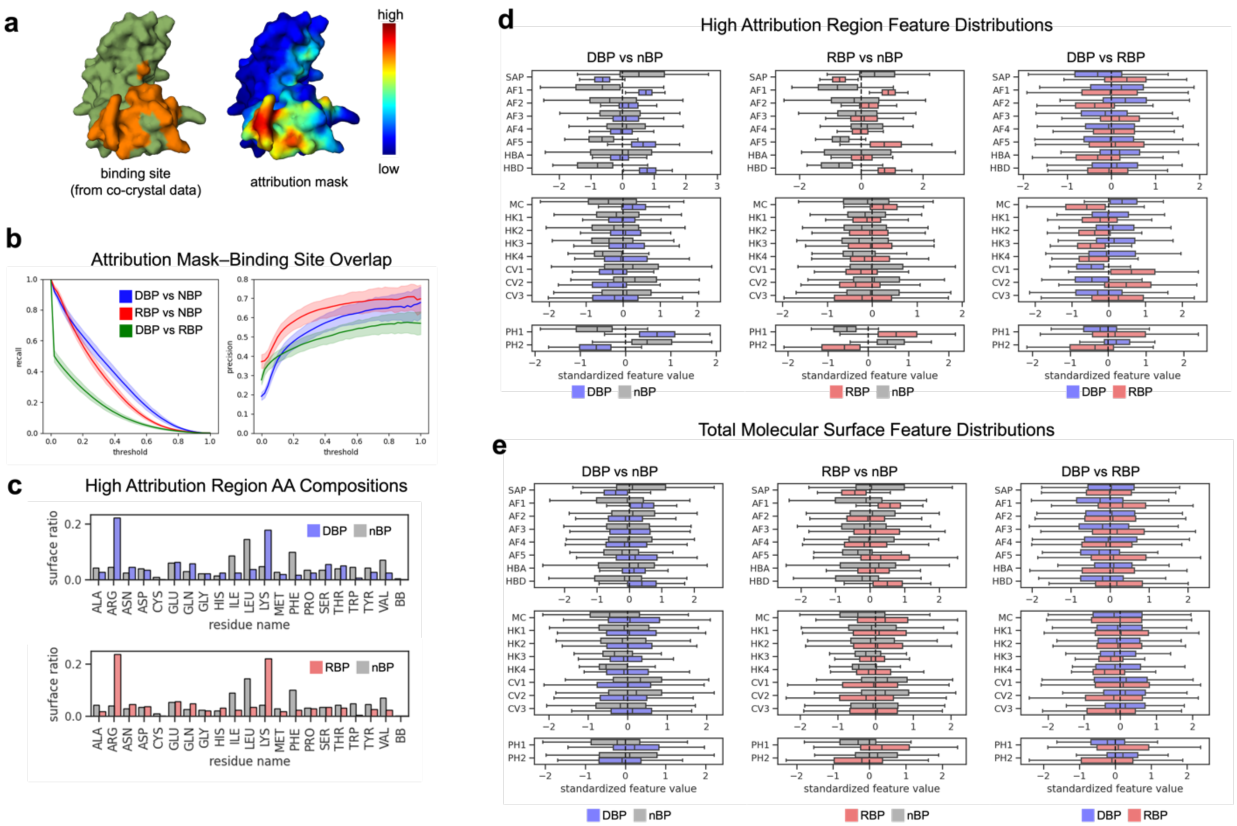


**Supplementary Figure 2: Spatial attribution comparisons and correlation with binding sites.** **a** Precision and recall curves computed from normalized spatial attribution masks and binding site labels transferred from proteins with experimental co-crystal structures available. **b** The proportion each residue side chain and the peptide backbone (BB) contributes to the total surface area within high spatial attribution regions. The top plot was computed from the DBP versus nBP model and the bottom plot from the RBP versus nBP model. **c** Feature distributions within high spatial attribution regions for each classification model. **d** Feature distributions over the entire molecular surface for each classification model. Much less separation is seen in the distributions relative to those in panel **c**.

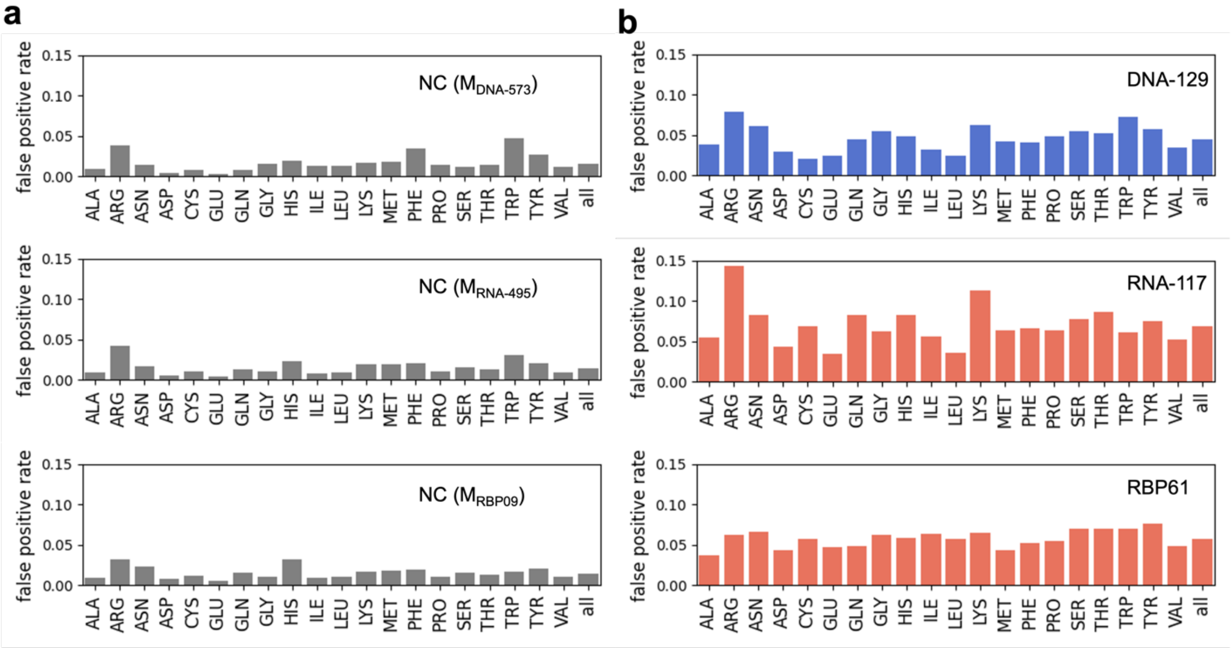


**Supplementary Figure 3: Residue–level false positive rates for binding site prediction.**
**a** False positive rates for each residue type from the negative control (NC) dataset and the three binding site prediction models ($M_{DNA-573}$, $M_{RNA-495}$ $M_{RBP09}$). **b** False positive rates for each residue type from the benchmark test sets.

**a**

Experimental Training Datasets

| Name | Type | # of target chains* | Positive Count | Negative Count |
|---|---|---|---|---|
| RBP09 | RBP | 808 | 33,160 | 208,931 |
| DNA-573 | DBP | 573 | 14,479 | 145,404 |
| RNA-495 | RBP | 495 | 14,609 | 122,290 |

Experimental Test Datasets

| Name | Type | # of target chains* | Positive Count | Negative Count |
|---|---|---|---|---|
| RBP61 | RBP | 61 | 2974 | 8729 |
| DNA-181 | DBP | 181 | 3208 | 72,050 |
| DNA-129 | DBP | 129 | 2240 | 35,275 |
| RNA-117 | RBP | 117 | 2031 | 35,314 |
| NC | NBP | 340 | 0 | 133,611 |

**b**

AlphaFold Datasets

| Name | Total Structures | Positive Count | Negative Count |
|---|---|---|---|
| DnP-6784 | 6784 | 3392 | 3392 |
| RnP-6046 | 6046 | 3392 | 2654 |
| RDP-6046 | 6046 | 2654 | 3392 |

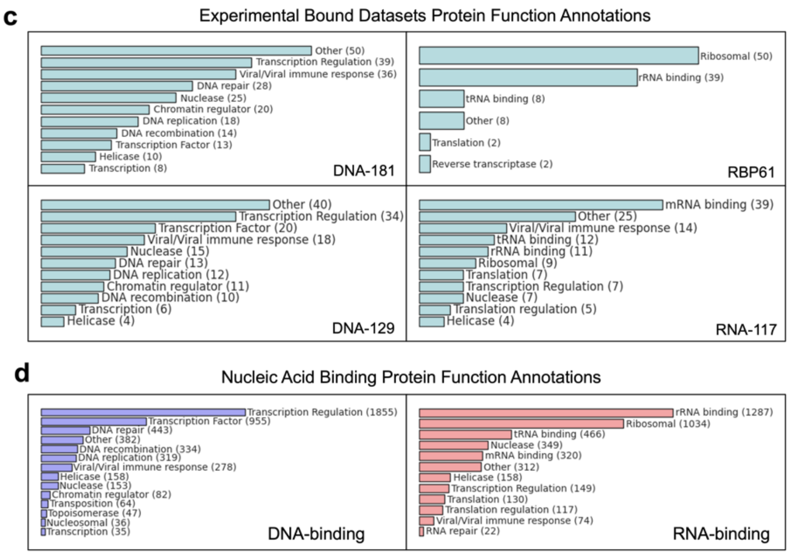

**Supplementary Figure 4: Statistics and functional annotations of datasets used in this study. a** Statistics for binding site prediction training and test sets containing experimentally determined protein structures. Positive count refers to the number of residues labeled as nucleic acid binding, and negative count is the number of solvent exposed residues that do not interaction with nucleic acids in the observed complex. **b** Statistics for binding function prediction datasets comprising protein structures predicted by AlphaFold2. Positive count refers to number of proteins labeled as the positive class (DNA binding for DnP-6784, RNA binding for RNB-6046 and RNA binding for RDP-6046) and negative count is the number of proteins labeled as the negative class. **c**–**d** Functional annotations derived from UniProt and Gene Ontology[1] molecular function annotations for proteins in the datasets shown in panels **a**–**b**.
